# Viral non-coding RNAs hijack host Pumilio proteins to regulate host transcripts

**DOI:** 10.1101/2025.11.27.691032

**Authors:** Nhi Phan, Paulina Pawlica

## Abstract

Viral non-coding RNAs (ncRNAs) often perform multiple functions during persistent infection, but how they interface with host RNA-binding proteins remains incompletely understood. Herpesvirus saimiri (HVS), a T-tropic γ-herpesvirus that induces aggressive lymphomas in primates, during its latency expresses abundant small nuclear RNAs HSUR1 and HSUR2. HSUR1 is known to trigger target-directed microRNA decay (TDMD) of host miR-27a, and HSUR2 can function as a miRNA-dependent adaptor; here we show that both HSURs also bind host Pumilio proteins (PUM1 and PUM2) via perfectly conserved Pumilio response elements (PREs). PUM proteins are potent post-transcriptional regulators that bind PREs usually located in 3′ UTRs of mRNAs to repress protein production. In Jurkat T cells, wild-type HSUR1/2, but not PRE-mutant variants, shift cells from a naïve-like toward an effector/memory-like state. This PRE-dependent shift is evidenced by increased surface expression of CXCR3, a chemokine receptor of activated effector T cells, and CD57, a marker of terminally differentiated T and NK cells. Our data support a model in which HSURs sequester a fraction of PUM proteins. Consistently, depletion of PUM1 or PUM2 phenocopies HSUR-induced changes, with more abundant PUM1 exerting the dominant effect. Consistently, depletion of PUM1 or PUM2 phenocopies HSUR-induced changes, with more abundant PUM1 exerting the dominant effect. In BJAB B cells, HSUR1 still drives PRE-dependent gene regulation, but the affected genes only partially overlap with those in Jurkat cells, and some transcripts even show reversed direction of change, revealing strong cell-type specificity. By identifying PUM1/2 as key host partners of HSUR ncRNAs, this work uncovers an underappreciated role for PUM proteins in shaping T- and B-cell states and reveals an additional way in which *γ*-herpesviral ncRNAs reprogram lymphocytes to promote viral persistence.

## Introduction

Viruses have evolved to maximize the information carried by their compact genomes, so when a viral non-coding RNA (ncRNA) is produced, it is functional and often fulfills multiple roles^1^. In particular, during programmed γ-herpesvirus latency, when only a limited subset of viral proteins is expressed, viral ncRNAs perform numerous functions. For example, pathogenic human γ-herpesviruses such as Epstein–Barr virus (EBV) and Kaposi sarcoma–associated herpesvirus (KSHV) rely heavily on ncRNAs: EBV, which is linked to ∼1% of human cancers^2^, establishes long-term latency in memory B cells, and in latency 0 the only viral genes expressed are ncRNAs, underscoring their central roles in cell transformation and viral persistence^3^. Latency-associated viral ncRNAs drive cell transformation and oncogenesis, yet many of their functions remain poorly understood^1,3–5^.

Here, we focus on ncRNAs from the classic γ-herpesvirus herpesvirus saimiri (HVS). HVS induces aggressive T-cell leukemias and lymphomas in New World primate species, such as tamarins, marmosets, and owl monkeys^6^, and some strains can transform human T cells^7^. During latency, HVS abundantly produces seven Sm-class small nuclear RNAs (snRNAs), known as HSUR1 through HSUR7 (HVS U-rich RNAs)^8–10^. Unlike host snRNAs, which mediate messenger RNA (mRNA) splicing, HSURs perform non-canonical and novel functions, providing valuable insights into host RNA biology^11–13^. For example, HSUR1 was the first *bona fide* RNA shown to induce the decay of a microRNA (miRNA). miRNAs are small ncRNAs that post-transcriptionally regulate gene expression by binding mRNAs and inhibiting their translation and/or promoting their degradation^14^. The observation that HSUR1 binds the host miRNA miR-27a and triggers its decay, rather than being regulated by it, led to the discovery of the process now known as target-directed miRNA degradation (TDMD)^11,15–18^. miR-27a regulates T cell activation and effector function, and its selective removal by HSUR1 promotes the persistence of virally transformed T cells^19^. HSUR2 selectively removes host pro-apoptotic mRNAs by serving as an adaptor between these mRNAs and the miRNA-induced silencing complex (miRISC)^13^. Finally, both HSUR1 and HSUR2 (as well as HSUR5) bind host miR-142-3p, which is necessary for these functions^12^.

Pumilio (PUM) proteins are sequence-specific RNA-binding proteins (RBPs) conserved across plants, fungi, and animals^20^. They play critical roles in growth, development, neurological function, and fertility, and mutations in PUM genes are associated with human diseases, including ataxia and leukemia^20,21^. As essential post-transcriptional regulators of gene expression, PUMs bind specific sequences known as Pumilio Response Elements (PREs), with the consensus 5′-UGUAHAUA (where H is A, C, or U), in mRNAs to control their stability and translation. Humans have two PUM paralogs, PUM1 and PUM2, which bind the same PRE consensus sequence, resulting in a highly similar mRNA repertoire regulated by each protein. However, despite this target similarity, PUM1 contains an additional N-terminal domain, and it is becoming increasingly clear that the functions of PUM1 and PUM2 do not completely overlap^22–27^.

In the current study, we show that HSUR1 and HSUR2, through their conserved PREs, bind the host RBPs PUM1 and PUM2. In a human T-cell line, we identify host genes that are upregulated by both HSURs in a PRE-dependent manner. Depleting PUM proteins demonstrates that these genes are normally repressed by PUMs, indicating that HSURs act by antagonizing PUM-mediated repression. The genes de-repressed by HSURs include regulators of T-cell specialization, revealing an underappreciated role for PUM proteins in shaping T-cell biology. Finally, in a B-cell context, HSUR1 exhibits PRE-dependent activities that only partially overlap with those in T cells, underscoring the cell-type specificity of PUM function.

## Results

### HSUR1 and HSUR2 bind host PUM proteins via conserved PREs

Despite their distinct activities^11,13^, HSUR1 and HSUR2 share several features, including binding sites for miR-142-3p, AU-rich elements (AREs) that mediate interaction with HuR/ELAVL1, Sm-binding sites, and a terminal stem–loop (**Fig. 1A**)^8,28^. Here, we uncover an additional shared feature—conserved PREs embedded in the loop of a structurally conserved hairpin in both HSUR1 and HSUR2 (**Fig. 1A, B**). Notably, unlike the miR-27a and miR-16 binding sites, these PREs are absolutely conserved across all HVS strains and in homologous ncRNAs from the closely related herpesvirus ateles (**Fig. S1A**), suggesting an important evolutionarily conserved function. To test whether HSURs interact with PUM proteins, we used marmoset T cells transformed with either wild-type (WT) HVS or a virus lacking HSUR1 and HSUR2 (Δ2A)^29^. RNA immunoprecipitation (RIP) of PUM1 and PUM2 showed that both HSURs are bound by both PUM proteins (**Fig. 1C**). PUM protein levels were comparable in both cell lines, indicating that HSURs do not alter PUM expression (**Fig. 1D**). To test whether the conserved PREs mediate PUM binding, we mutated the PREs in HSUR1 and HSUR2 and expressed these variants, along with an empty-vector (EV) control, in the human T-cell line Jurkat (**Fig. 1E**), which we selected because HVS is T-tropic. PRE mutations did not affect ncRNA expression. HSUR1 retained the ability to trigger miR-27a degradation, although this activity was partially reduced (**Figs. 1E, S1B**). As expected, neither ncRNA altered PUM protein levels (**Fig. S1C**). Consistently, WT HSURs, but not PRE-mutant variants (Pm), bound PUM proteins (**Fig. 1F**). Together, these results demonstrate that HSURs bind host PUM proteins via conserved PREs, pointing to yet another function for these viral ncRNAs.

**Figure 1.**
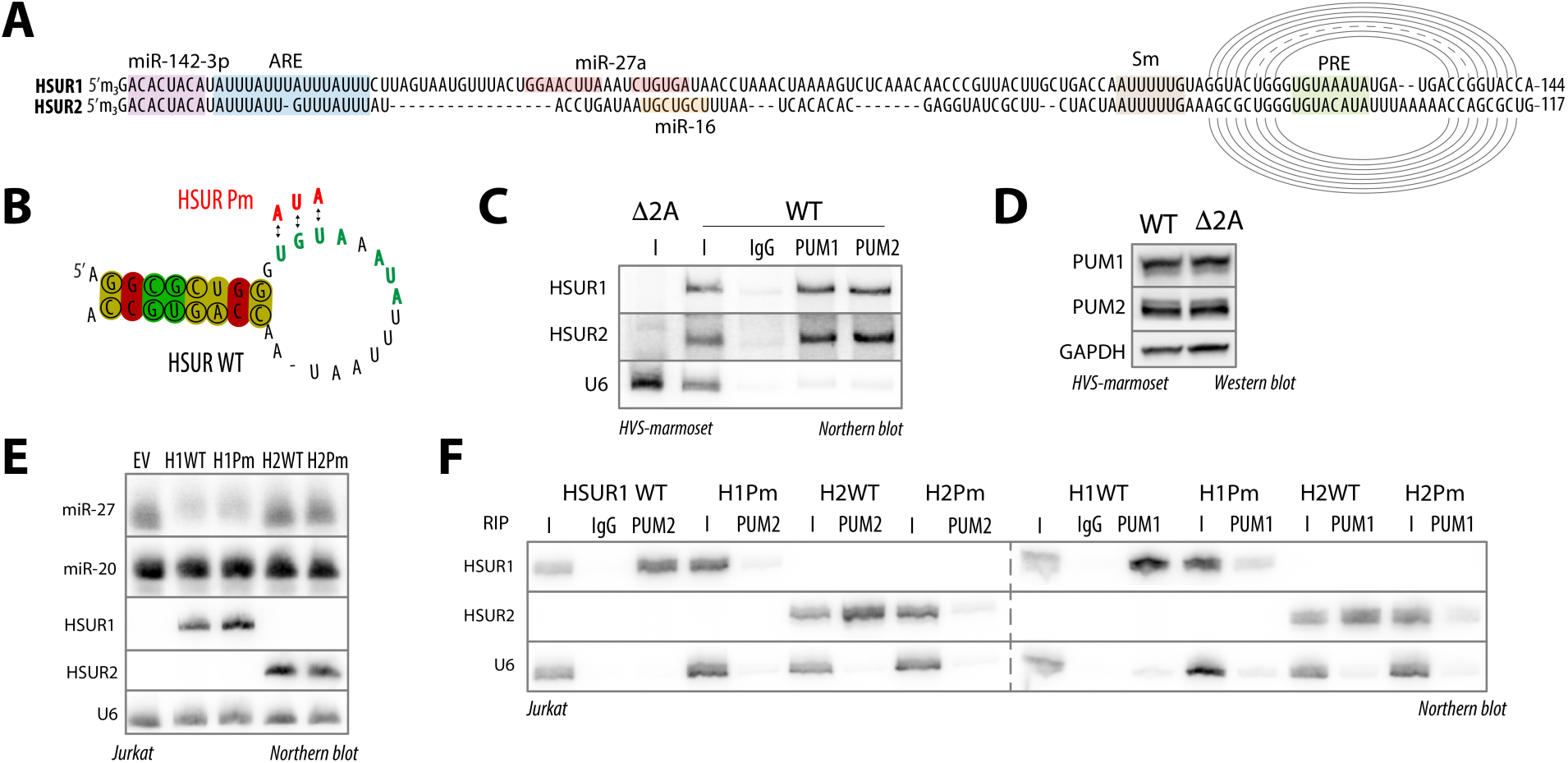
HSUR1 and HSUR2 bind host PUM proteins via conserved PREs. **(A)** HSUR1 and HSUR2 contain conserved PREs. Alignment of HSUR1 and HSUR2 indicating binding sites for miRNAs and RNA-binding proteins. Arches indicate base pairing. ARE, AU-rich element; Sm, binding site for Sm proteins; PRE, Pumilio response element. **(B)** The 3′-terminal stem–loop of HSUR1 and HSUR2 is structurally conserved across multiple HVS isolates and related herpesvirus ateles. Common secondary structure was predicted by RNAalifold^56^. Circles indicate consistent and covarying nucleotides. Red indicates base pairs with no sequence variation; ochre and green indicate base pairs with two and three different types of base pairs, respectively. Mutations in the PRE used in this study are indicated. **(C)** PUM1 and PUM2 bind both HSUR1 and HSUR2. RNA immunoprecipitation from marmoset T cells transformed with WT HVS using anti-PUM1 antibody, anti-PUM2 antibody, or IgG control. I, input (5%). Input from cells transduced with virus lacking HSUR1 and HSUR2 (Δ2A) was included as an additional negative control. **(D)** PUM1 and PUM2 levels are not affected by the presence of HSUR1 and HSUR2. Western blot analysis of protein levels in marmoset cells transformed with either WT or Δ2A HVS. (E) PRE mutations do not affect HSUR expression. Representative Northern blot of RNAs from Jurkat cells transduced with the indicated HSUR variants or EV, probed for ncRNAs. **(F)** HSURs bind PUM proteins via PREs. RNA immunoprecipitation from Jurkat cells transduced with the indicated HSUR variants using anti-PUM1 antibody, anti-PUM2 antibody, or IgG control. I, input (5%).

### HSUR1 and HSUR2 regulate multiple genes in a PRE-dependent manner

To understand why HSURs bind PUM proteins, we performed RNA sequencing (RNA-seq) of Jurkat cells stably transduced with WT HSUR1 or HSUR2 or their respective Pm variants, alongside the EV control. We investigated HSUR1 and HSUR2 simultaneously to strengthen our conclusion about their PRE-dependent impact on the host transcriptome. Consistent with their distinct functions, many dysregulated genes were unique to either HSUR1 or HSUR2; however, both ncRNAs also affected a substantial set of overlapping genes (**Fig. 1, Fig. S2A**). Since HSUR1 triggers TDMD of miR-27a, we assessed the mRNA levels of miR-27a targets (from miRTarBase^30^) and found that they were slightly (2.6%) but significantly higher than non-targets, consistent with their global de-repression (**Fig. S2B**). PRE mutation reduced this effect, consistent with a contribution of PUM binding to HSUR1-mediated TDMD. In contrast, we did not observe the previously described HSUR2-mediated effect on anti-apoptotic genes in this system. Comparison of changes in transcript levels for each ncRNA variant relative to EV revealed many genes whose upregulation required an intact PRE, including multiple genes commonly affected by HSUR1 and HSUR2 WT, but not by their Pm variants (**Fig. 2A**). Direct comparison of each WT HSUR to its Pm mutant further identified numerous genes upregulated in a PRE-dependent fashion (**Fig. 2B**). Because PUM binding most often destabilizes its targets^20^, we hypothesized that increased expression of these genes reflects reduced PUM activity. For subsequent analyses, we defined PUM targets as transcripts previously reported to associate with PUM proteins^31–33^ and containing at least one conserved PRE in the 3′ UTR. Among mRNAs dysregulated by HSURs in a PRE-dependent manner, PUM targets comprised 30% for HSUR1 and 35% for HSUR2, compared with 21% transcriptome-wide. Similarly, a global analysis of PUM targets revealed a modest but statistically significant increase in their levels, 2.2% and 1.9% for HSUR1 and HSUR2, respectively, and PRE mutation markedly decreased this effect (**Figs. 2C, S2C**). These findings are consistent with a model in which HSURs interfere with PUM-mediated repression, and we speculate that the remaining altered genes are indirect and reflect downstream regulation by primary PUM targets. In summary, we identified host mRNAs that are upregulated by both HSURs in a PRE-dependent manner, and our data suggest that this upregulation stems from partial inhibition of PUM function.

**Figure 2.**
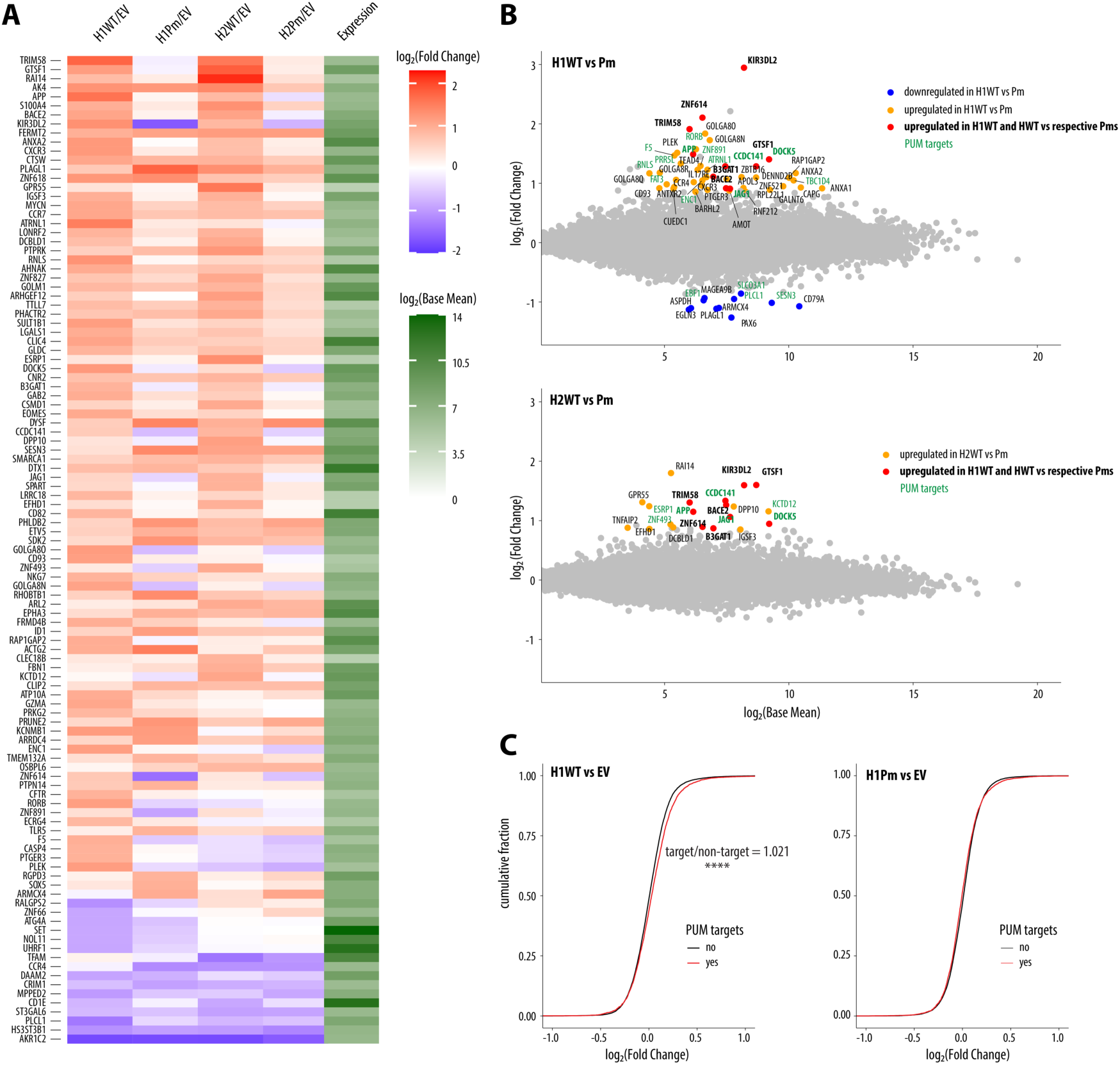
HSUR1 and HSUR2 regulate multiple genes in a PRE-dependent manner. **(A)** WT HSURs, but not their PRE-mutated counterparts (Pm), upregulate many common genes in Jurkat cells. Heatmap shows log₂FC values for transcripts significantly dysregulated by either HSUR variant relative to EV. Differentially expressed genes were defined as those with |log₂FC| ≥ 0.85 and adjusted p-value ≤ 0.05 (edgeR^57^; n = 3). (B) HSURs upregulate overlapping genes in a PRE-dependent manner. Scatter plots compare WT HSURs with their Pm mutants; genes upregulated for both HSURs are bolded, and PUM targets (defend as bound by PUM proteins^31–33^ and containing at least one conserved PRE in the 3′ UTR) are shown in green. (C) HSUR1 WT, but not its PRE mutant, causes a global increase in PUM-target mRNA levels, as shown by cumulative distribution plots of log₂FC for PUM targets versus non-targets (****, Wilcoxon rank-sum test, p ≤ 0.0001). FC, fold change.

### HSUR1 and HSUR2 act by counteracting PUM-mediated repression

Our RNA-seq data revealed that HSURs impact PUM targets in a PRE-dependent manner, yet most dysregulated mRNAs are not classified as PUM targets. In our analysis defining PUM targets, we focused on canonical PREs, even though PUMs have also been shown to bind shorter, non-canonical sequences^33,34^. To determine whether genes upregulated by HSURs in a PRE-dependent manner are PUM targets, we performed anti-PUM1 and anti-PUM2 RIP. As expected, the lncRNA NORAD, which has ∼18 PREs^35^, was strongly bound by both PUMs, and other PUM targets such as *JAG1* (Jagged1) and *APP* (amyloid precursor protein) were also enriched (**Fig. 3A**). In contrast, the “non-target” HSUR-upregulated genes *GTSF1* (gametocyte-specific factor 1), *KIR3DL2* (killer cell immunoglobulin-like receptor 3DL2), and *TRIM58* (tripartite motif–containing 58) showed minimal association with PUM proteins. To ask whether these genes, despite not being detectably bound by PUMs, are nonetheless regulated by PUM proteins, we generated polyclonal knockouts of either PUM1 or PUM2 in Jurkat cells. *PUM1* mRNA levels were reduced 2.8-fold (±0.66), which correlated with a 2.6-fold (±0.96) increase in *PUM2* mRNA levels, consistent with auto- and cross-regulation of PUM production via PREs within their own mRNAs (Fig. 3B)^20,32^. *PUM2* mRNA levels were decreased 2.3-fold (±0.39) in PUM2-depleted cells. PUM1 depletion had a larger impact on transcript levels than PUM2 depletion, likely reflecting its higher abundance and previously noted distinct functional role. Using previously reported estimates of PUM copy number in HCT116 cells^36^ as a reference, we estimated that in Jurkat cells PUM1 is ∼6-fold more abundant than PUM2 (**Fig. S3A**). Upon depletion of PUM proteins, we observed an increase in the same genes that are upregulated by HSURs in a PRE-dependent manner, including “non-targets” such as *GTSF1* and *KIR3DL2*, supporting the idea that the changes observed in the presence of HSURs are driven by altered PUM activity (**Fig. 3B**). To test whether expression of WT HSURs decreases PUM binding to endogenous targets, we performed RIP with anti-PUM antibodies in Jurkat cells expressing HSUR variants and assessed NORAD association by RT–qPCR. Although not statistically significant, we observed a trend toward reduced NORAD enrichment in the presence of WT HSURs (**Fig. S3B**). We next examined whether PUM targets regulated by HSURs in a PRE-dependent manner share specific features and found that lower-abundance mRNAs were slightly more susceptible to this regulation (**Fig. S3C**). In summary, we show that genes upregulated by HSURs in a PRE-dependent manner are regulated by PUM proteins, supporting the notion that HSURs can de-repress a subset of PUM targets.

**Figure 3.**
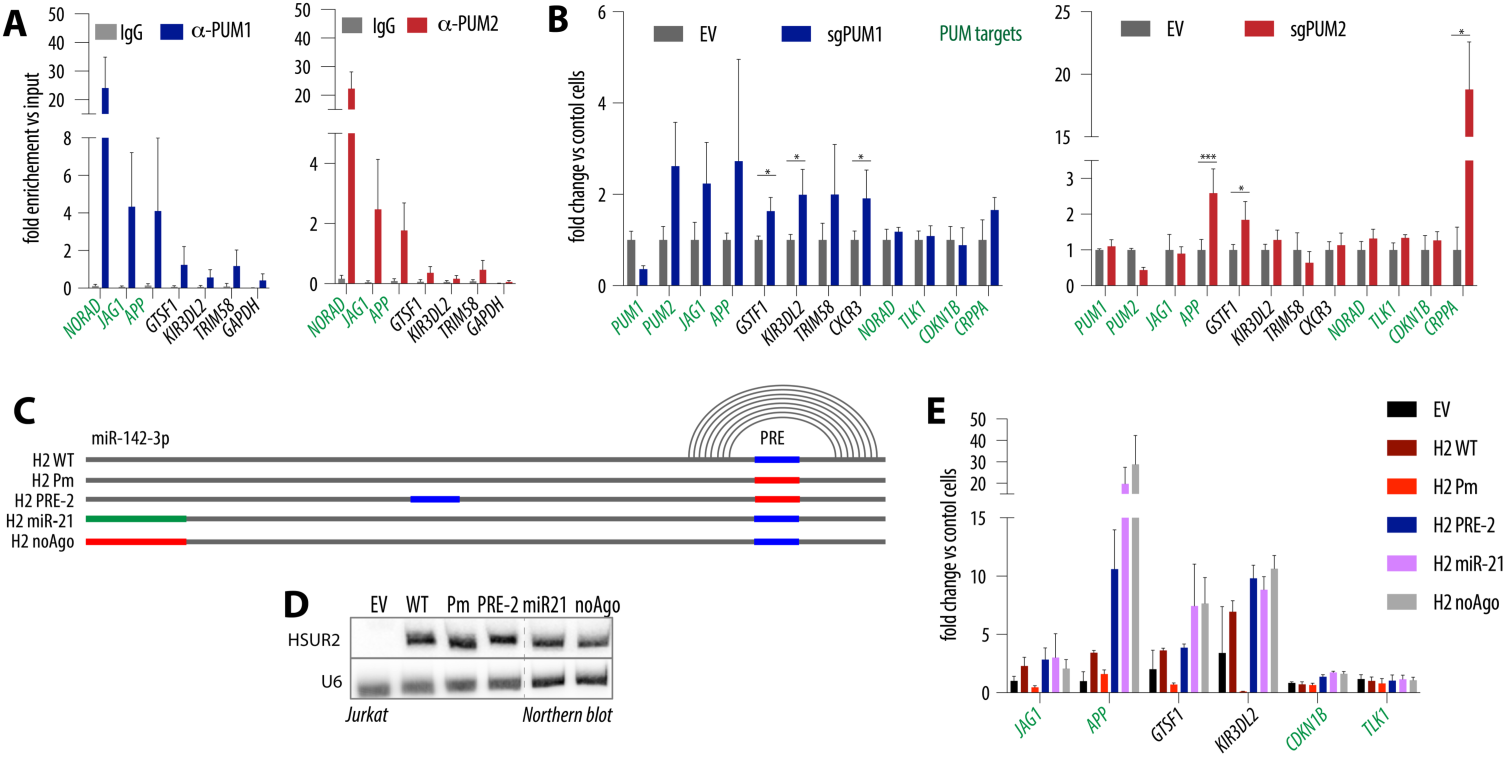
HSUR1 and HSUR2 act by counteracting PUM-mediated repression. **(A)** Only a subset of genes upregulated by HSURs or PUM loss are direct PUM targets. RIP in Jurkat cells using anti-PUM1 (left), anti-PUM2 (right), or IgG control, shown as enrichment over input; mean of 3 independent experiments ± SD. **(B)** Genes upregulated by HSURs are also induced upon PUM depletion. RNA levels of selected transcripts in polyclonal PUM1 (left) or PUM2 (right) knockouts in Jurkat cells; mean of 3 independent experiments ± SD; *p ≤ 0.05, ***p ≤ 0.001 (paired t-test). **(C)** Schematic of HSUR2 mutants designed to identify additional elements involved in regulation of PUM-dependent targets. **(D)** Northern blot showing expression of HSUR2 variants in Jurkat cells. **(E)** Neither PRE location nor miR-142-3p binding affects HSUR2’s ability to increase PUM-regulated transcripts. Relative mRNA levels of selected transcripts in Jurkat cells expressing the indicated HSUR2 variants; mean of 3 independent experiments ± SD.

### HSUR2 increases PUM-regulated transcripts regardless of PRE location or miR-142-3p binding

To define which HSURs elements are required for regulation of this common gene set, we performed mutagenesis of HSUR2, chosen because of its smaller size. To test whether the PRE must reside in the conserved 3′ stem-loop, we reintroduced a PRE at an alternative position in the H2 Pm variant (**Figs. 3C and S3D**). Because miR-142-3p binding is essential for other HSUR functions^12^, we either swapped this site for miR-21 (H2 miR21), as done previously, or generated a mutant that is not predicted to bind any miRNA (H2 noAgo). We left the Sm-binding site and terminal stem–loop intact, as they are required for snRNA assembly and stability. These variants were stably expressed in Jurkat cells (**Fig. 3D**). Notably, none of the mutations impaired the ability of PRE-containing HSUR2 to upregulate the identified genes, and in some cases gene induction exceeded that of HSUR2 WT (**Fig. 3E**). In summary, neither PRE location nor miR-142-3p binding affects HSUR2’s ability to increase transcripts regulated by PUM in this system.

### HSURs upregulate PUM-dependent genes that shape T-cell state

To understand the biological significance of PREs in HSURs, we used GSEA^37,38^ to identify biological processes regulated by HSURs in a PRE-dependent manner in Jurkat cells and found enrichment of pathways related to cell adhesion, migration, and T cell activation (**Fig. 4A**). When we focused specifically on T cell–related gene sets, PRE-containing HSURs were associated with reduced naïve-like CD4⁺ T cell signatures and a shift toward memory-like transcriptional programs (**Fig. S4A**). This prompted us to examine genes regulated by HSUR in a PRE-dependent way encoding surface receptors involved in T cell specialization. CXCR3 is a chemokine receptor induced on effector T cells and NK cells upon activation^39^. B3GAT1 encodes a β1,3-glucuronyltransferase that generates the HNK-1/CD57 epitope, a marker of terminally differentiated NK cells and subsets of CD8⁺ and CD4⁺ T cells^40^. Using surface staining, we validated that HSURs increase CXCR3 and CD57 expression in a PRE-dependent manner (**Fig. 4B–D**). Moreover, HSUR1 functionally promoted an effector-like state, as PMA stimulation led to increased surface CD107a (LAMP-1), a standard degranulation marker^41^, in a PRE-dependent manner (**Fig. S4B,C**). To test whether these effects are mediated by PUM regulation, we performed the same surface staining in Jurkat cells with polyclonal knockouts of either PUM1 or PUM2. Depletion of both PUMs led to a marked increase in CD57, but only PUM1 loss increased CXCR3, whereas PUM2 depletion reduced CXCR3 levels (**Fig. 4E**). These results support the idea that genes upregulated by HSURs in a PRE-dependent manner reflect reduced PUM activity, and that PUM1 and PUM2 are not fully redundant but have at least partially distinct functions^22–27^. In summary, our results demonstrate that HSURs regulate PUM-dependent genes encoding key surface receptors, promoting a transition of Jurkat T cells from a naïve-like to an activated, effector- and memory-like state. These findings highlight a role for PUM proteins in controlling T-cell state and effector function.

**Figure 4.**
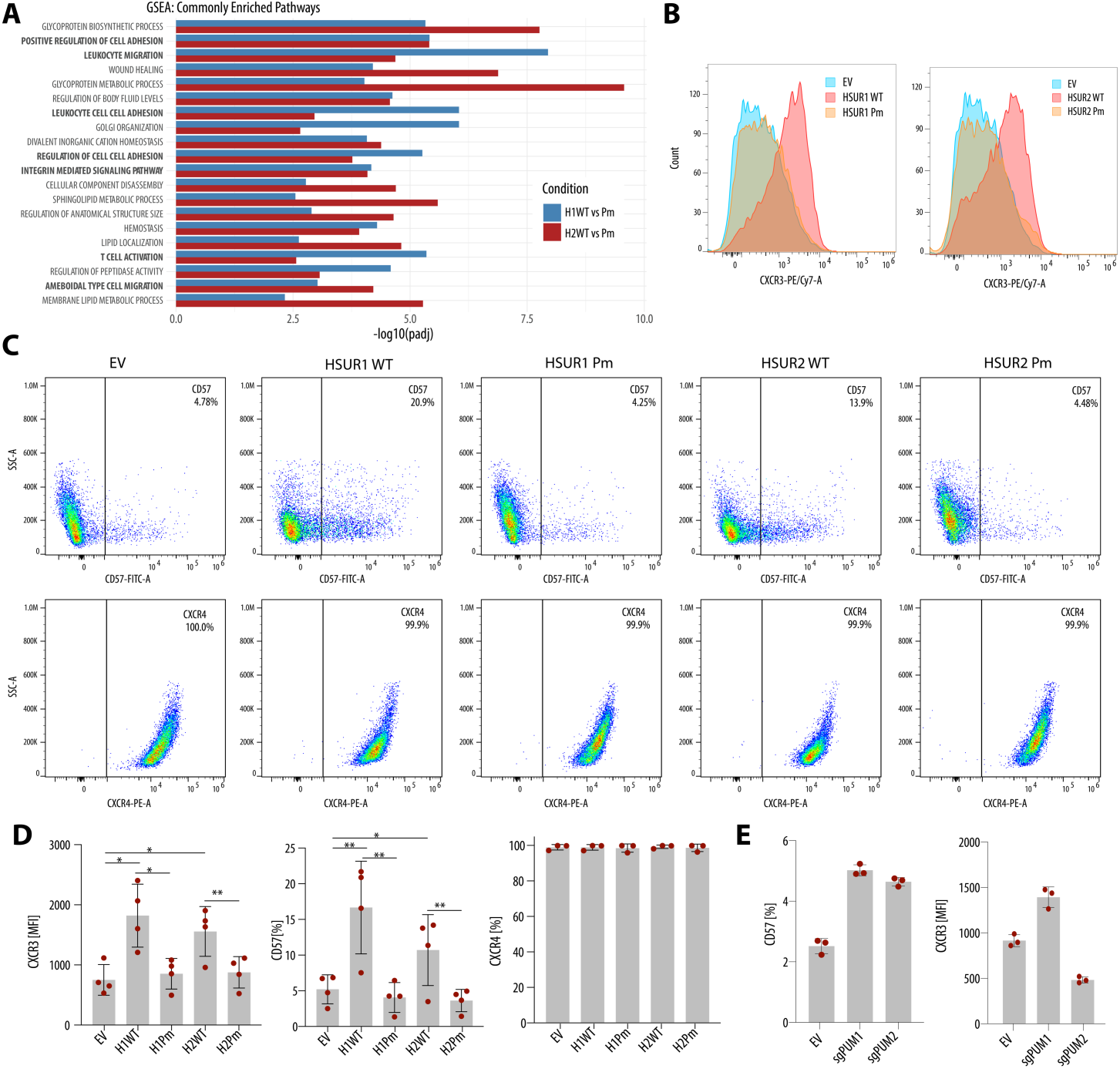
HSURs upregulate PUM-dependent genes that shape T-cell state. **(A)** Gene set enrichment analysis (GSEA) using GO gene sets from MSigDB^38^ showing biological processes altered by WT HSUR1 and HSUR2 as compared to their respective Pm mutants. **(B)** WT HSURs increase CXCR3 surface expression. Representative MFI histograms of CXCR3 staining in Jurkat cells expressing the indicated HSUR variants. **(C)** WT HSURs increase CD57 levels on the cell surface, whereas CXCR4 remains unchanged. Representative flow cytometry plots of CD57 or CXCR4 staining in Jurkat cells expressing the indicated HSUR variants. **(D)** Quantification of the data in **(B)** and **(C)** from 4 independent experiments (mean ± SD). **(F)** Impact of polyclonal PUM1 or PUM2 knockout on CXCR3 and CD57 surface levels. Data are from a triplicate experiment (mean ± SD). EV, empty vector; MFI, mean fluorescence intensity; GO, Gene Ontology.

### PRE-dependent HSUR regulation reveals that PUMs have different roles in different cell types

To test whether PRE-dependent HSUR regulation of PUM targets is conserved across cell types, we expressed HSUR1 WT and its Pm variant in the B-cell line BJAB. As in Jurkat cells, PRE mutation did not alter HSUR1 abundance and only modestly reduced its ability to trigger TDMD of miR-27a (**Fig. S5**). RNA-seq of BJAB cells expressing HSUR1 WT or HSUR1 Pm revealed many significantly dysregulated genes, although the magnitude was more modest than in Jurkat cells (Fig. 5A). This difference may reflect altered PUM activity or target usage in B cells. Consistent with this idea, only a small subset of genes altered in Jurkat cells was also changed in BJAB cells. Interestingly, *PAX6* mRNA decreases in a PRE-dependent manner in Jurkat cells but increases in BJAB cells. As in Jurkat cells, many HSUR1-regulated transcripts in BJAB are not classified as PUM targets and likely represent indirect, downstream effects of altered PUM-target regulation. Nevertheless, cumulative distribution analysis showed a mild global shift toward higher expression of PUM targets in HSUR1 WT compared with the Pm variant (**Fig. 5B**), indicating that HSUR1 still acts, at least in part, through PUM in B cells. In summary, these data suggest that PRE-dependent HSUR regulation operates in both T and B cells but engages distinct sets of PUM targets, underscoring cell type–specific roles for PUM proteins.

**Figure 5.**
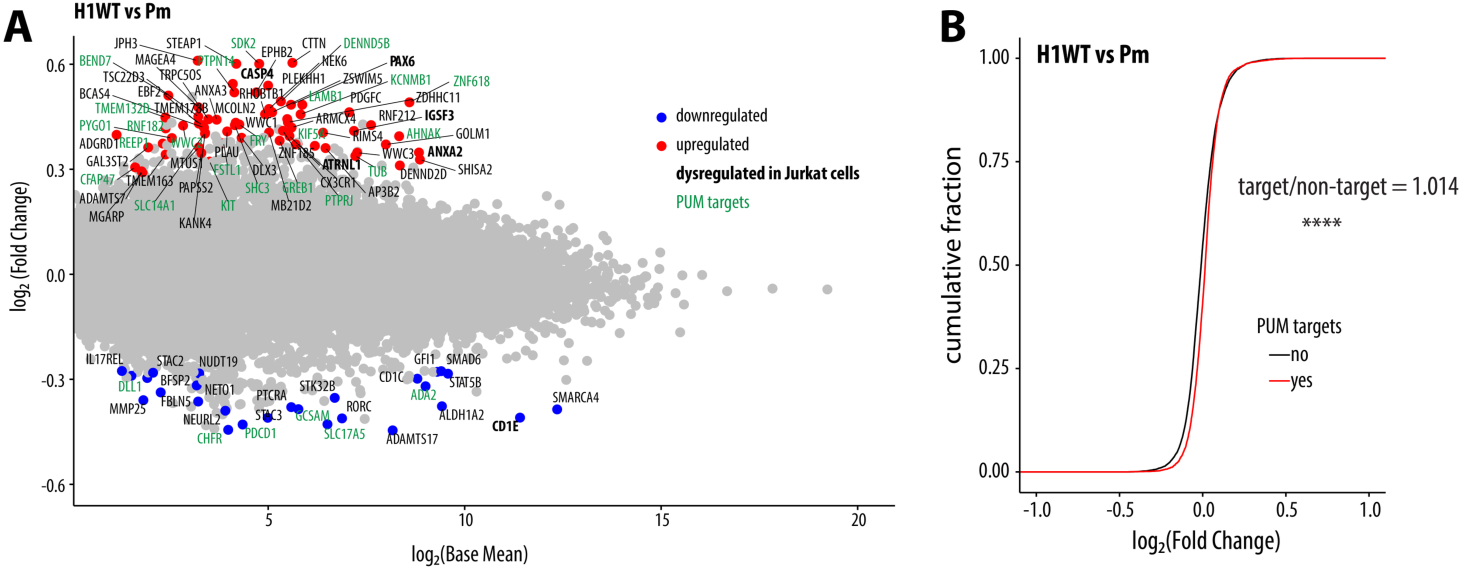
PRE-dependent HSUR regulation reveals that PUMs have different roles in different cell types. (A) HSUR1 WT upregulates multiple transcripts in a PRE-dependent manner in BJAB cells. Differentially expressed genes were defined as those with |log₂FC| ≥ 0.26 and p-value ≤ 0.05 (edgeR, REF; n = 3). Genes also changed in Jurkat cells are bolded, and PUM targets are marked in green. (B) HSUR1 WT, as compared to the Pm variant, leads to a global increase in PUM targets, as shown by cumulative distribution plots of log₂FC for PUM targets vs non-targets (****, Wilcoxon rank-sum test, p ≤ 0.0001). FC – fold change.

## Discussion

Here, we discover that HSUR1 and HSUR2 bind PUM1 and PUM2, allowing these viral ncRNAs to modulate the PUM program. We show that HSUR expression partially relieves PUM-mediated repression. Mechanistically, our data are consistent with a sponging mechanism in which HSURs sequester a fraction of PUM proteins, most likely PUM1, given its higher abundance, away from endogenous targets, with low-abundance PUM targets being slightly more sensitive to this perturbation. Thus, HSURs, although likely less potent, resemble the host lncRNA NORAD, which titrates PUM proteins^36,42^. Although global changes in PUM targets are modest (typically 2–3%), PRE-dependent upregulation of genes such as *CXCR3*, *KIR3DL2*, and *B3GAT1*/CD57 results in functionally meaningful reprogramming of T-cell state.

Our findings underscore the context dependence of PUM biology. PUM1 and PUM2 are enriched in hematopoietic stem and progenitor cells and contribute to blood cell development, yet their roles in mature lymphocytes remain largely unknown^43^. Their overexpression promotes expansion of stem and leukemia cells, highlighting how PUM dysregulation can drive hematologic disease^44,45^. HSUR1-induced PUM dysregulation differs between T and B cells, as only a subset of genes dysregulated by HSUR1 in a PRE-dependent manner overlaps between Jurkat and BJAB cells, and genes such as *PAX6* show opposite PRE-dependent behavior in the two contexts. Moreover, our data corroborate previous reports that PUM1 and PUM2 are not fully redundant^22–27^; CXCR3 regulation is an example in which loss of PUM1 and PUM2 has distinct, opposing effects. Together, these observations argue that PUM regulation is highly cell type–specific and that viral ncRNAs can serve as useful tools to probe this layer of control.

In the context of *γ*-herpesvirus latency, shifting T cells toward more activated, longer-lived, metabolically active states is likely advantageous, creating a stable niche that can sustain persistent infection and, over time, may contribute to oncogenic risk. A second consequence of modulating PUM activity may relate to previously reported antiviral functions of PUM1^26,46,47^, raising the possibility that HSUR–PUM interactions both rewire host gene expression and blunt intrinsic antiviral activity of PUM1. HSURs are therefore not single-function effectors but multipronged regulators of host RNA metabolism. HSUR1 drives TDMD of miR-27a, HSUR2 can serve as an adaptor for miRNA-guided repression, and we now show that both ncRNAs additionally modulate PUM-dependent repression via conserved PREs^11,13^. This combination of miRNA decay, miRNA-guided silencing, and RBP titration could give HVS unusually fine control over T-cell fate and survival. PREs also appear to enhance the TDMD activity of HSUR1 by a mechanism that remains unclear. One possibility is that cytoplasmic PUM proteins prolong the cytoplasmic phase of HSUR biogenesis, where Ago-loaded miRNA complexes reside and TDMD occurs.

In summary, by identifying PUM1/2 as key host partners of multifunctional HSUR ncRNAs, we uncover an underappreciated role for PUM proteins in shaping T- and B-cell transcriptional programs. HSUR–PUM interactions give γ-herpesviruses a way to subtly reprogram lymphocytes toward effector/memory-like states, likely promoting viral persistence.

## Acknowledgements

This work was funded by NIH grants R35 GM150649 (P.P.) and S10OD026880 and S10OD030463 (High-Performance Computing at Mount Sinai). Additional support was provided by the Clinical and Translational Science Awards (CTSA) grant UL1 TR004419 from the National Center for Advancing Translational Sciences. We are grateful to the members of the Jean Lim lab for advice and support with flow cytometry. We thank the members of the Mount Sinai Flow Cytometry Core, especially Alessandro Marins dos Santos. We thank Drs. Anan-Lena Steckelberg and Ben Kleaveland for helpful discussions, and Yelizaveta Zaytseva for critically reading the manuscript.

## Author Contributions

N.P. and P.P. designed, performed and interpreted experiments. P.P. wrote the manuscript.

## Declaration of interests

Authors declare no competing interests.

## Data availability

Sequenced reads will be deposited in the NCBI Gene Expression Omnibus (GEO) database.

## Materials and methods

### Cells

HEK293T (ATCC) and HCT116 (ATCC) cells were grown at 37°C with 5% CO₂ in DMEM containing 10% fetal bovine serum (FBS, HyClone), 2 mM L-glutamine (GIBCO), and 100 I.U./mL penicillin and 100 μg/mL streptomycin (Pen/Strep, GIBCO). Jurkat and BJAB cells (kind gifts from Joan Steitz) were grown in RPMI with 10% FBS, 2 mM L-glutamine, and Pen/Strep. Marmoset (Callithrix jacchus) T cells transformed with either WT or Δ2A HVS^48^ were grown in RPMI with 20% FBS, 10 μg/mL BME, 2 mM L-glutamine, and Pen/Strep. The cell lines exhibit expected morphology and are mycoplasma-negative.

### Generation of stable cell lines expressing HSUR variants or sgRNAs targeting PUM proteins

Plasmids generated previously^16,49^ were subjected to QuikChange mutagenesis. sgRNAs in the lentiCRISPRv2 (Addgene #49535) backbone targeting PUM proteins were described previously. Lentiviral vectors were prepared in HEK293T cells by transfecting the backbones together with packaging plasmids pMD.G2 and psPAX2 (Addgene #12259, #12260). Jurkat and BJAB cells were transduced with lentiviral vectors at high multiplicities of infection in the presence of 8 μg/mL polybrene and spinoculation, followed by puromycin (GIBCO) selection. For polyclonal PUM knockouts, two guides per gene were used.

### RNA Immunoprecipitation (RIP)

RIP was performed as previously described^26^. Briefly, 5×10⁶ Jurkat or marmoset cells were lysed in 1 ml of NET-2 buffer (50 mM Tris [pH 7.5], 150 mM NaCl, and 0.05% Nonidet P-40) in the presence of RNase inhibitor (NEB) and Protease Inhibitor Cocktail (Pierce). The lysate was divided into 3 parts and each was incubated with 30 µl Dynabeads Protein G (Invitrogen) coupled to 10 µg of antibodies: anti-PUM1 (Bethyl), anti-PUM2 (Bethyl), and an IgG control (Sigma). After IP, the beads were washed 7 times in NET-2 buffer. Beads were resuspended in TRIzol (Invitrogen), and RNA was extracted and analyzed by Northern blot and RT-qPCR.

### Northern blot analysis

10-15 µg of total RNA was separated by 15% urea-PAGE, electro-transferred to Hybond-NX membrane (Amersham) and crosslinked either with UV (conditions) or with 1-ethyl-3-(3-dimethylaminopropyl) carbodiimide (EDC)^50^ when miRNAs were assayed. RNAs were detected using ^32^P 5’-radiolabeled DNA probes (see Table 1), by storage phosphor screens and Typhoon. Densitometry was performed by using Fiji.

**Table 1.**
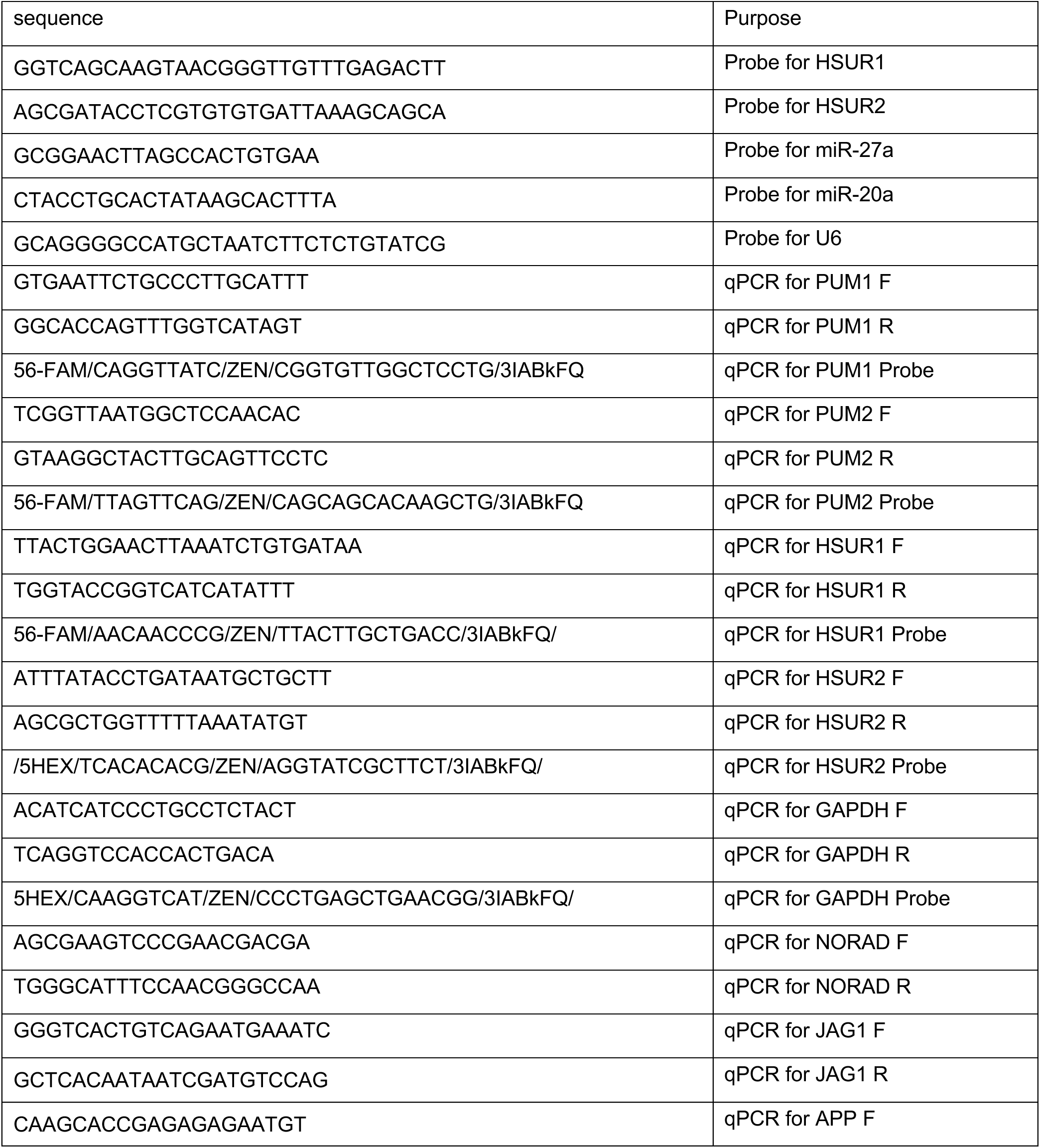
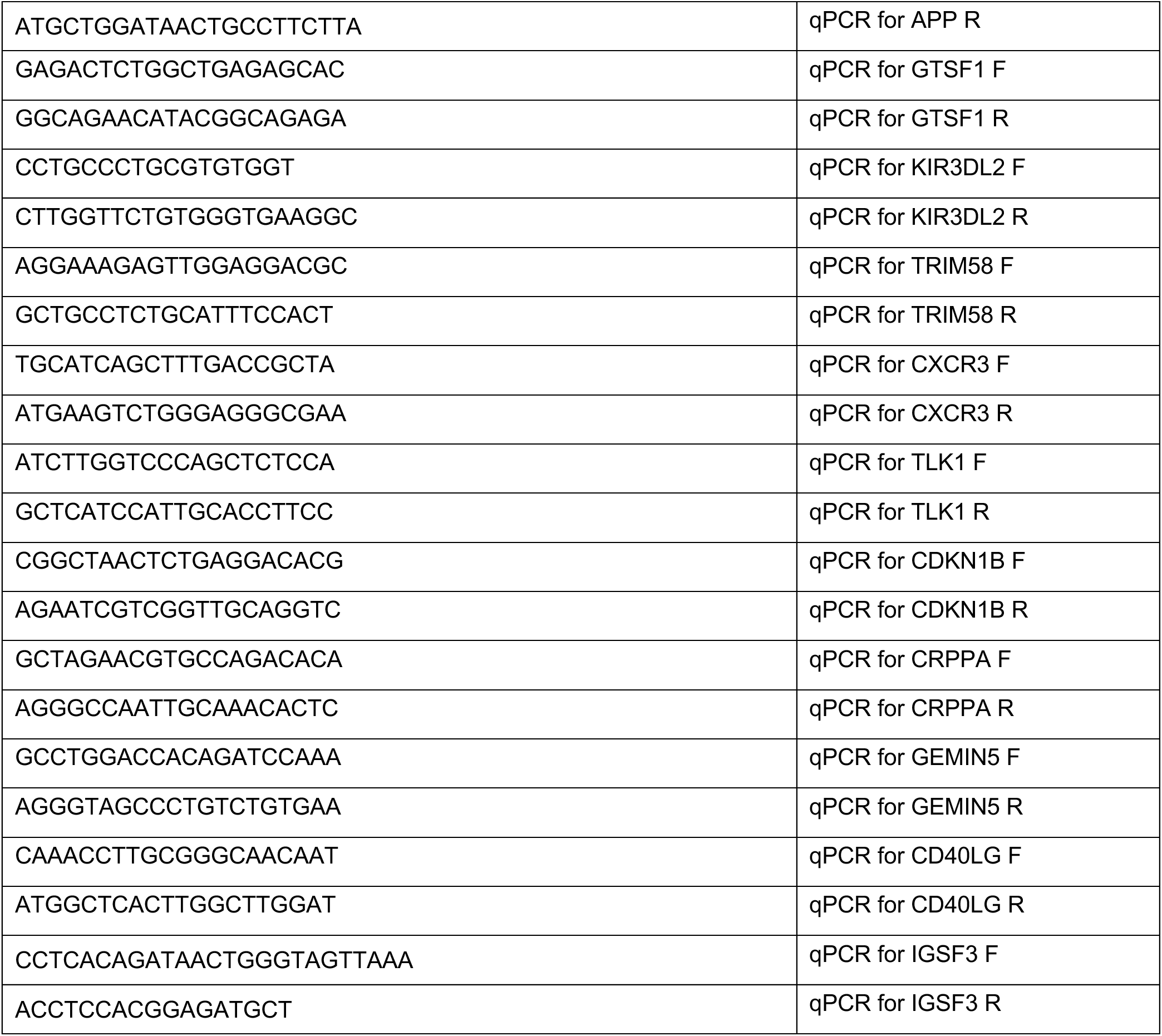

### Western blot analysis

Cells were lysed in IP Lysis buffer (Pierce) containing protease inhibitor cocktails (Pierce). 10-15 µg of total protein was separated on a gradient SDS-PAGE gel (Invitrogen), and electro-transferred to a PVDF membrane (BioRad). After blocking with 5% milk in TBST (20 mM Tris [pH 7.5], 150 mM NaCl, 0.1% Tween 20), the membrane was probed with the appropriate antibodies and detected using ECL Plus Western Blotting Substrate (Pierce) on a ChemiDoc MP Imaging System. Primary antibodies used were anti-PUM1 (Abcam), anti-PUM2 (Bethyl) and anti-GAPDH (Sigma).

### Reverse transcriptase qPCR

Total RNA was extracted from TRIzol according to the manufacturer’s recommendations. 1ug RNA was reverse transcribed using ProtoScript II and oligo(dT). Quantitative PCR was performed using either SYBR Green qPCR Master Mix (Thermo Fisher) or Luna qPCR Master Mix (NEB). The reactions were performed according to the manufacturer’s instructions. Sequences are listed in Table1.

### RNA sequencing and sequencing analysis

RNA sequencing and analysis. RNA-seq libraries were prepared using the NEBNext Ultra II Directional RNA Library Kit (NEB #E7760) with either the rRNA depletion module (NEB #E7400) or the poly(A) mRNA magnetic isolation module (NEB #E7490), according to the manufacturer’s instructions. Libraries were amplified for 9 cycles and sequenced on Illumina NovaSeq platforms (Azenta). Reads were mapped with STAR^51^ and genes were counted with featureCounts^52^. Differential expression was determined using DESeq2 with normal shrinkage^53^. GSEA was analyzed using fgsea^37^ and gene sets from MSigDB^38^. PUM targets were defined as transcripts (1) previously reported to bind PUM proteins in high-throughput experiments^32,33,54^ and (2) containing consensus PREs in their 3′ UTRs. miR-27a targets were taken from miRTarBase^55^, excluding “weak” targets.

### Cell surface staining

3 × 10⁵ cells were washed with PBS and stained for viability using the LIVE/DEAD™ Fixable Violet Dead Cell Stain Kit (Thermo Fisher) for 10 min at room temperature. Cells were then incubated on ice for 30 min with the following antibodies: FITC-conjugated anti-human CD57 (HNK-1; BioLegend), PE/Cyanine7-conjugated anti-human CD183 (CXCR3) (G025H7; BioLegend), anti-human CD107a (LAMP-1) (H4A3; BioLegend), and PE-conjugated anti-human CD184 (CXCR4) (12G5; BioLegend). After three washes with FACS buffer, cells were fixed with 2% paraformaldehyde. Flow cytometry was performed on an Attune NxT Flow Cytometer (Thermo Fisher), and data were analyzed using FlowJo software (version 8.8.4; Tree Star, Ashland, OR, USA).

---

**Figure S1.**
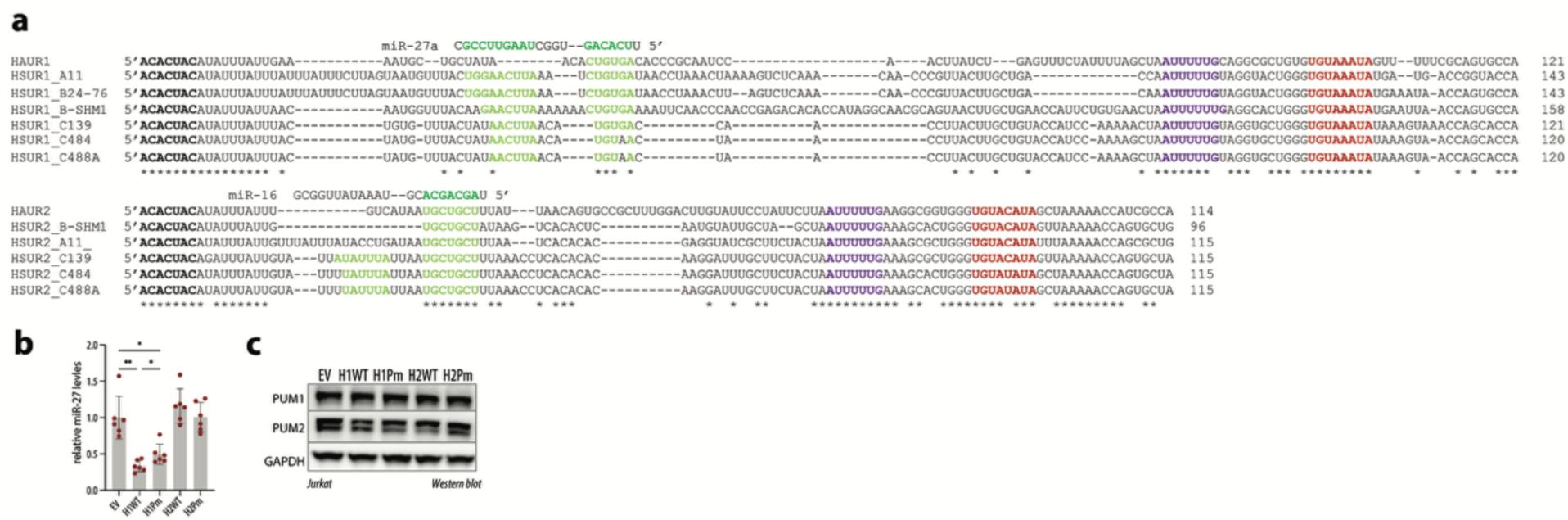
Related to Fig. 1. **(A)** PREs are conserved in HSUR1 and HSUR2 across multiple HVS isolates and related herpesvirus ateles. **(B)** Quantification of miR-27a levels normalized to miR-20; *p ≤ 0.05 and **p ≤ 0.01 by paired t-test. **(C)** PUM1 and PUM2 levels are not affected by the presence of HSUR variants. Western blot analysis of protein levels in Jurkat cells transduced with the indicated variants.

**Figure S2.**
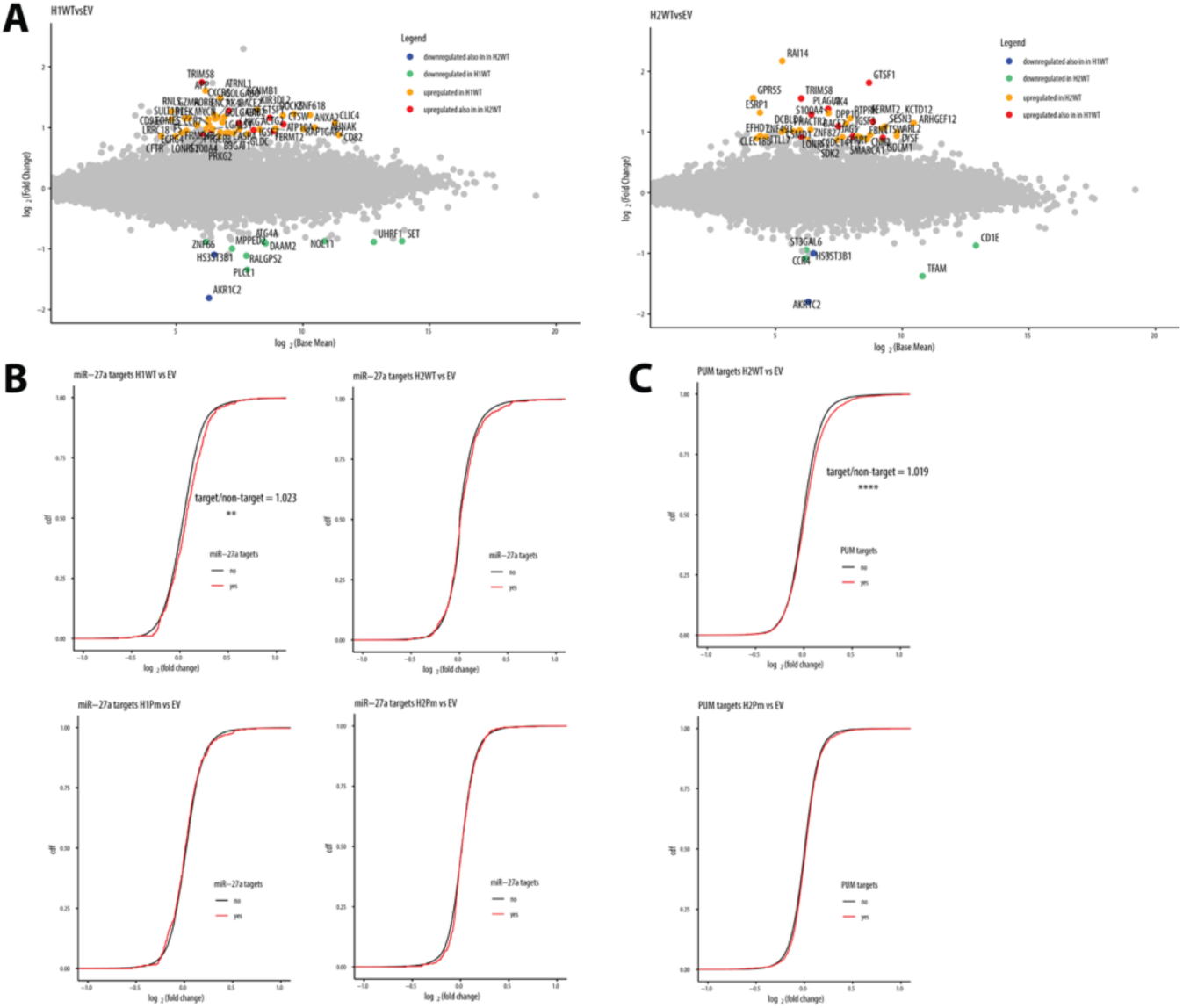
Related to Fig. 2. **(A**) Impact of WT HSURs on RNA levels in Jurkat cells. Shown are transcripts changing at least 2-fold with an adjusted p-value ≤ 0.05 (n = 3). **(B)** miR-27a targets exhibit a global increase in the presence of HSUR1, but not the other HSURs, as shown by cumulative distribution plots of log₂FC for miR-27a targets (from TarBase^55^) versus non-targets (**, p-value from Wilcoxon test). **(C)** HSUR2 WT leads to a global stabilization of PUM targets, as shown by cumulative distribution plots of log₂FC for PUM targets versus non-targets (****, p-value from Wilcoxon test). FC fold change.

**Figure S3.**
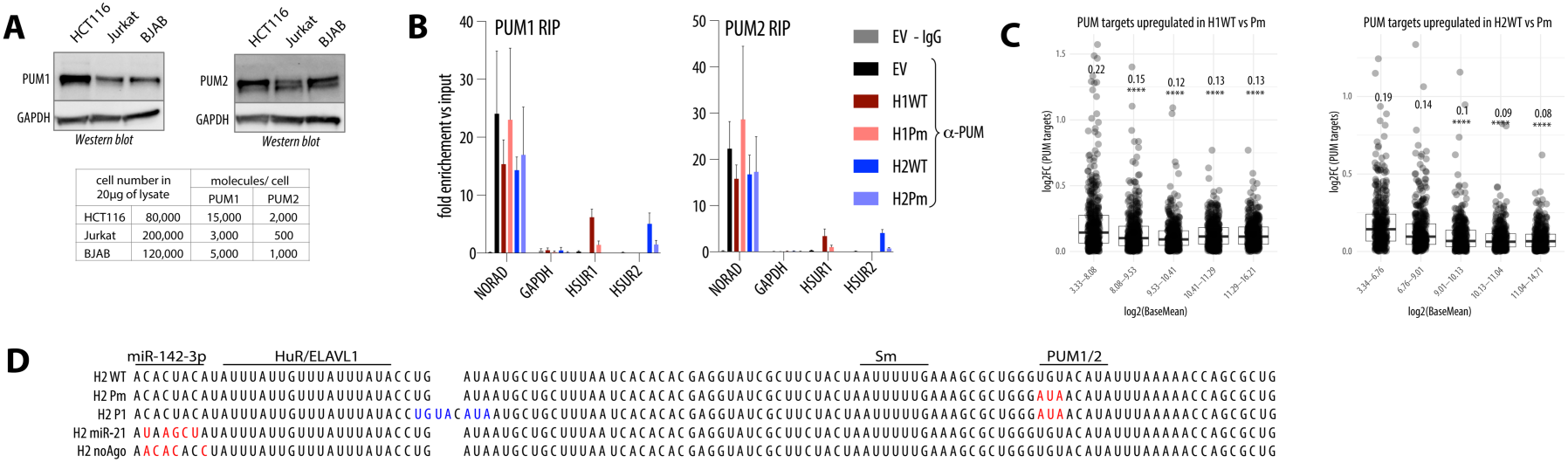
Related to Fig. 3. **(A**) Estimated numbers of PUM1 and PUM2 protein molecules per Jurkat cell relative to published values for HCT116 cells. **(B)** RNA immunoprecipitation in Jurkat cells using anti-PUM1 (left), anti-PUM2 (right), or IgG control, shown as enrichment over input; mean of 3 independent experiments ± SD. **(C)** Less abundant PUM targets are more strongly regulated by HSURs in a PRE-dependent manner; PUM targets were binned by expression level and PRE-dependent changes in expression are shown for each bin. **(D)** Schematic of HSUR2 mutants designed to identify additional elements involved in regulation of PUM-dependent targets.

**Figure S4.**
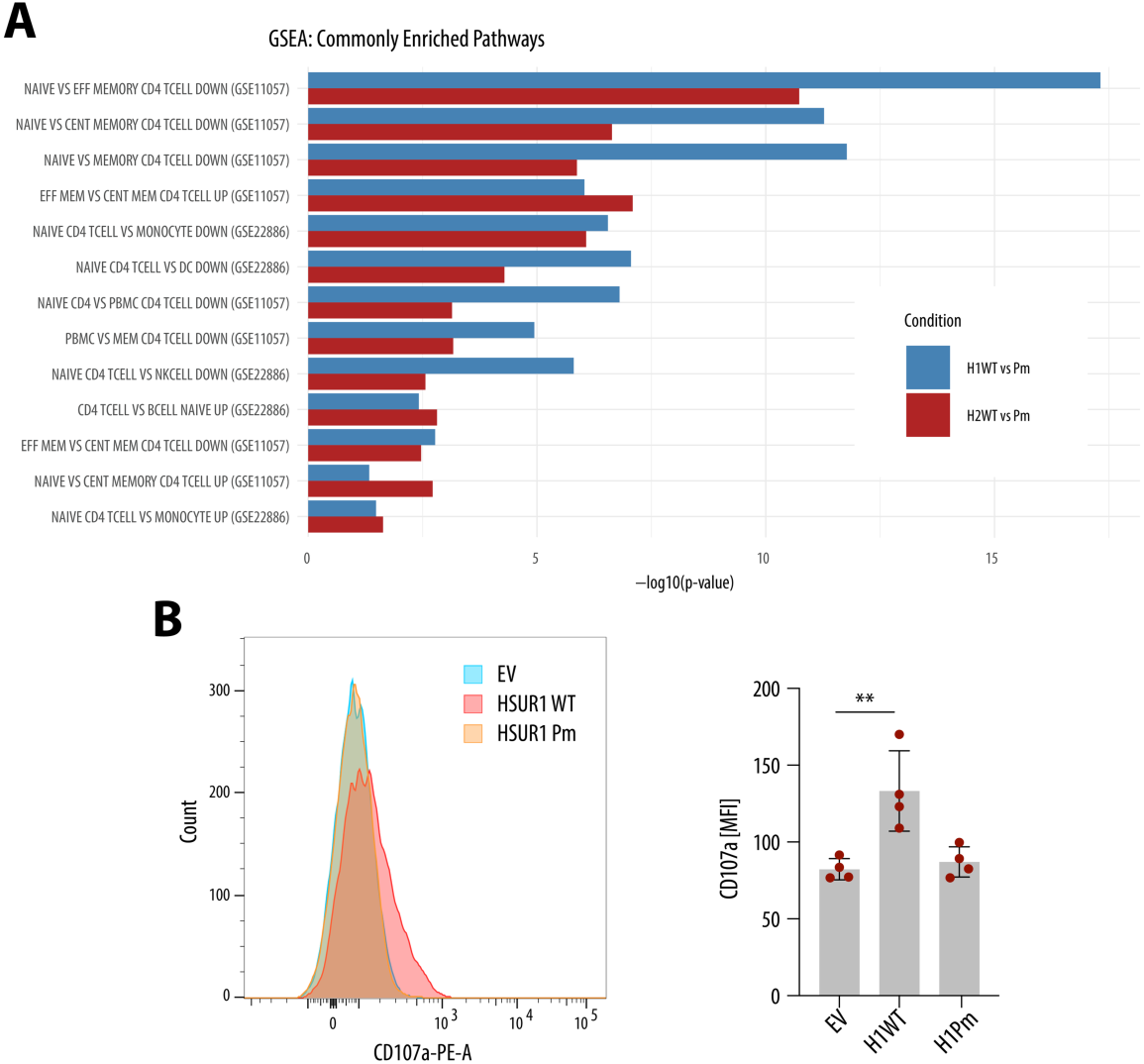
Related to Fig. 4. **(A)** Figure S4. Related to Fig. 4. (A) GSEA was restricted to MSigDB immunologic signatures containing “TCELL” in the gene set name and, for CD4⁺-focused analyses, further filtered to sets containing “CD4”; shown are gene sets altered by both WT HSUR1 and HSUR2 compared with their respective Pm mutants. **(B)** WT HSUR1 increases CD107a surface levels upon PMA stimulation; a representative plot and quantification of CD107a MFI from three independent experiments are shown (mean ± SD; p ≤ 0.01, paired t-test). MFI, mean fluorescence intensity.

**Figure S5.**
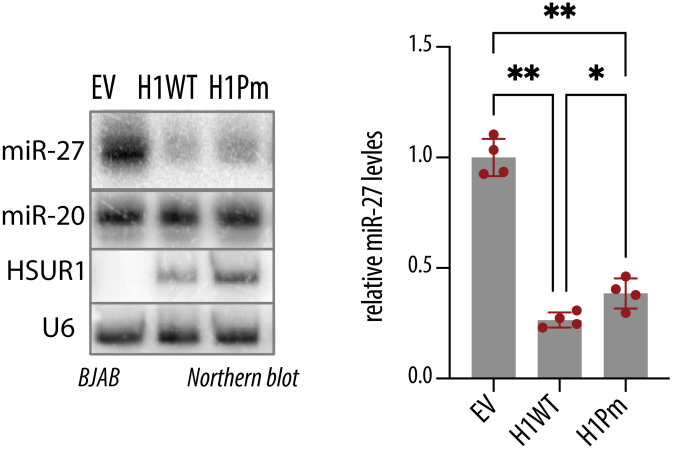
Related to Fig. 5. PRE mutation does not affect HSUR1 levels but slightly decreases TDMD of miR-27a. Representative Northern blot of RNAs from BJAB cells transduced with the indicated HSUR1 variants or empty vector (EV), probed for ncRNAs.

## Notes

### Competing Interest Statement

The authors have declared no competing interest.

